# The Diguanylate Cyclase YfiN of *Pseudomonas aeruginosa* Regulates Biofilm Maintenance in Response to Peroxide

**DOI:** 10.1101/2021.07.29.454409

**Authors:** Stefan Katharios-Lanwermeyer, Sophia A. Koval, Kaitlyn E. Barrack, G.A. O’Toole

## Abstract

*Pseudomonas aeruginosa* forms surface-attached communities that persist in the face of antimicrobial agents and environmental perturbation. Published work has found extracellular polysaccharide (EPS) production, regulation of motility and induction of stress response pathways as contributing to biofilm tolerance during such insults. However, little is known regarding the mechanism(s) whereby biofilm maintenance is regulated when exposed to such environmental challenges. Here, we provide evidence that the diguanylate cyclase YfiN is important for the regulation of biofilm maintenance when exposed to peroxide. We find that, compared to the wild type (WT), static biofilms of the Δ*yfiN* mutant exhibit a maintenance defect, which can be further exacerbated by exposure to peroxide (H_2_O_2_); this defect can be rescued through genetic complementation. Additionally, we found that the Δ*yfiN* mutant biofilms produce less c-di-GMP than WT, and that H_2_O_2_ treatment enhanced motility of surface-associated bacteria and increased cell death for the Δ*yfiN* mutant grown as a biofilm compared to WT biofilms. These data provide evidence that YfiN is required for biofilm maintenance by *P. aeruginosa*, via c-di-GMP signaling, to limit motility and protect viability in response to peroxide stress. These findings add to the growing recognition that biofilm maintenance by *P. aeruginosa* is an actively regulated process that is controlled, at least in part, by the wide array of c-di-GMP metabolizing enzymes found in this microbe.

**Importance:** We build on previous findings that suggest that *P. aeruginosa* utilizes c-di-GMP metabolizing enzymes to actively maintain a mature biofilm. Here, we explore how the diguanylate cyclase YfiN contributes to the regulation of biofilm maintenance during peroxide exposure. We find that mature *P. aeruginosa* biofilms require YfiN to synthesize c-di-GMP, regulate motility and to insure viability during peroxide stress. These findings provide further evidence that the modulation of c-di-GMP in response to environmental signals is an important mechanism by which biofilms are maintained.

## Introduction

*Pseudomonas aeruginosa* is a Gram-negative opportunistic pathogen that is found in settings as varied as natural aquatic environments, implanted catheters and the airways of patients afflicted with cystic fibrosis (1–3). *P. aeruginosa* forms robust biofilms, multicellular surface-attached aggregates that are recalcitrant to antimicrobial therapy and environmental insults (4, 5). For motile organisms such as *P. aeruginosa*, biofilms form through a multistep process that requires motility and motility appendages, surface-sensing and the production of extracellular polysaccharides (EPS) (6–9). Flagella mediate initial polar attachment of a cell to a surface, while pili facilitate irreversible surface attachment and microcolony formation (10, 11). After surface attachment, the PilY1 protein is secreted to the cell surface where it contributes to surface sensing and the activation of diguanylate cyclase SadC via a Type IV pili (TFP) dependent, outside in signal transduction pathway (8, 12). The activation of SadC results in increased bis-(3′,5′)-cyclic dimeric guanosine monophosphate (c-di-GMP) (12), a second messenger that coordinates surface behavior, virulence and the production of EPS (13–16). Concentrations of c-di-GMP are controlled enzymatically by diguanylate cyclases that synthesize this molecule and phosphodiesterases that degrade this second messenger (17–19). c-di-GMP controls events in biofilm formation beyond surface attachment. For example, once attached to a surface, the production of EPS, a process also regulated by c-di-GMP, provides microcolonies with architectural strength for continued growth and protection against predation and environmental challenges, such as antimicrobials, oxidative and osmotic stressors (4, 20–24).

Biofilms provide protection from a number of insults, including peroxide stress. For example, *Acinetobacter oleivorans* and *Helicobacter pylori* (25, 26) both respond to peroxides with increased EPS production and enhanced biofilm biomass, while for *P. aeruginosa* the peroxide (H_2_O_2_)-responsive OxyR regulates metabolism through the control of iron homeostasis and protein synthesis (27). However, little is known regarding how mature biofilms respond to the exposure to peroxides, including H_2_O_2_, an important issue given that molecules with such chemistry are produced by the immune system (3, 28, 29).

While early surface attachment events and the processes that drive biofilm maturation (i.e., reversible/irreversible attachment, microcolony formation, EPS production) have received a great deal of attention in the literature, less is known about the regulatory components needed for biofilm maintenance. In previous work in a collaborative effort with the Howell group, we found that RmcA and MorA, phosphodiesterases that break down c-di-GMP, regulate the maintenance of nutrient-limited biofilms (30). To assess if genes that encode diguanylate cyclase activity also contribute to biofilm maintenance, we utilized a mutant library containing clean deletions of all known single-domain DGCs (31). We screened mutants that individually lacked these DGC domains in a static assay to identify deficits in biofilm maintenance. Through this process, we identified YfiN (32) as a putative biofilm maintenance protein. The *yfiN* gene has previously been found to be a member of the *yfiBNR* operon (33). Repression of YfiN activity is mediated by its regulator, YfiR. YfiR activity is repressed when bound by YfiB under some conditions (33, 34). Oxidative stress is thought to play an important role in the regulation of YfiBNR activity, with the crystal structures of YfiR indicating denaturation of this protein in oxidizing conditions (35) Here, we investigate the role of YfiN in *P. aeruginosa* motility and biofilm maintenance, and how peroxide stress impacts these biological functions.

## Results

### The Δ*yfiN* mutant is defective for biofilm maintenance and susceptible to peroxide stress

We previously reported the use of a library of in-frame deletion mutants targeting each of ∼40 genes of *P. aeruginosa* PA14 predicted to participate in c-di-GMP synthesis or degradation (31). We used this library to determine if any of the genes that encode diguanylate cyclases (DGCs) contribute to the ability of *P. aeruginosa* to maintain a biofilm once the community has been established. To test this idea, strains carrying mutations in each of the DGCs of *P. aeruginosa* PA14 were grown statically in 96-well plates for 12h and 48h in M63 biofilm medium (**Table S1)**. As we have described previously, the identification of candidate mutants involved in biofilm maintenance is apparent through a biofilm defect at 48h, but not at 12h (30). From our initial screen of these DGC mutants, we found that that the strain carrying a mutation in the *yfiN* gene exhibited a biofilm comparable to WT at 12h (**Fig. 1A**, left panel) but displayed a significant defect at 48h, with lower amounts of biofilm biomass at this time point (**Fig. 1A**, middle panel).

**Figure 1.**
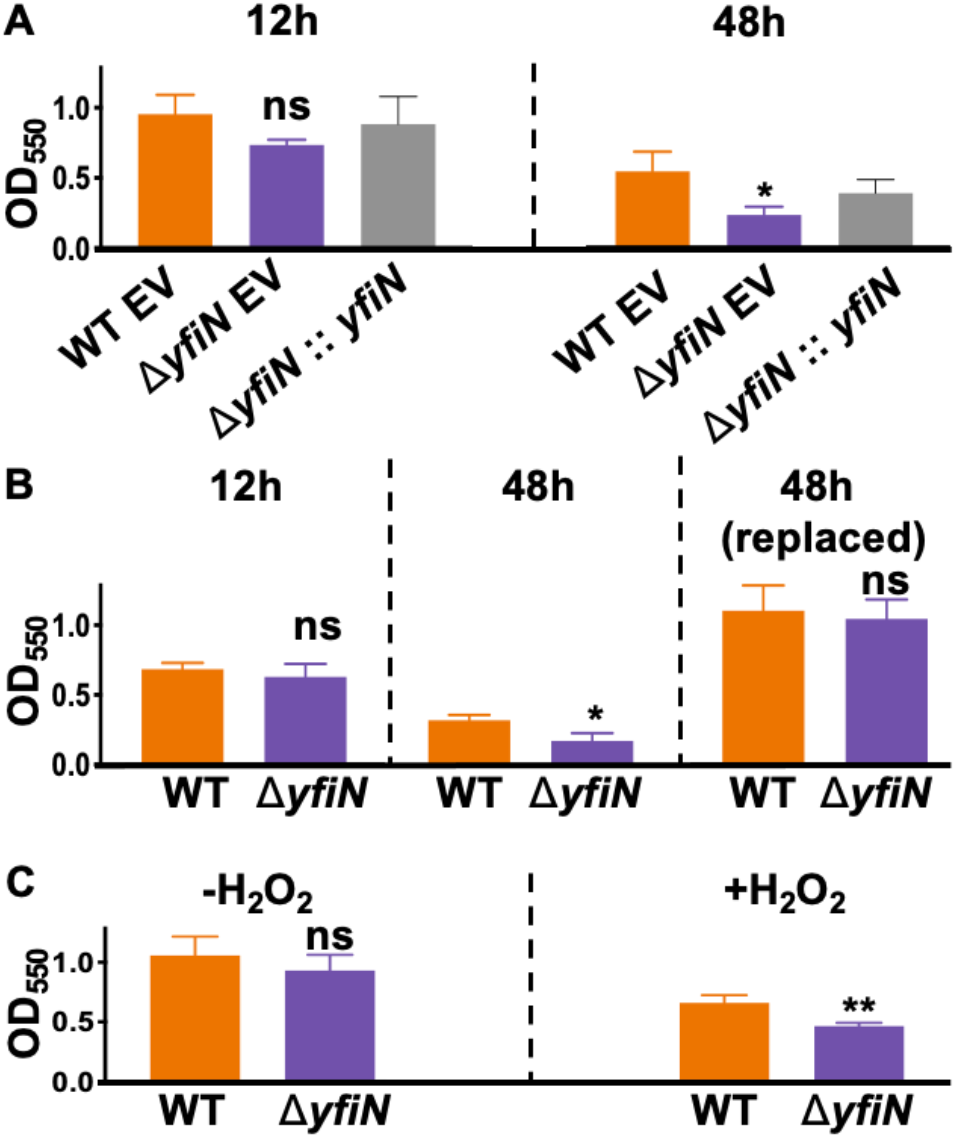
The Δ*yfiN* mutant is defective for biofilm maintenance and a pre-formed biofilm is reduced when exposed to peroxide. A) Biofilm formation by the WT, the WT carrying the pMQ70 empty vector (EV), the Δ*yfiN* mutant carrying the EV, and Δ*yfiN* mutant complemented with pMQ70-*yfiN* after 12h and 48h (left and right panel, respectively). The biofilm formed was assessed by staining with crystal violet (CV), the CV-stained biofilm solubilized in 30% glacial acetic acid and the absorbance (OD_550_) measured. B) Biofilm formation by WT and the Δ*yfiN* mutant was assessed in a 96-well plate assay. The strains were grown on M63 medium supplemented with arginine (0.4% w/v) and the biofilm measured after 12h (left panel), 48h (middle panel), or after 48h wherein the medium was replaced every 12h (right panel). The quantification of the biofilm by CV was performed as described in panel A. C) Biofilms of WT and the Δ*yfiN* mutant were grown for 12h, after which medium was replaced with control growth medium (left panel) or medium supplemented with 50 mM H_2_O_2_ (+H_2_O_2_) for an additional 12h of growth. Biofilm formation was assessed as described in panel A. For all panels, error bars represent standard deviation of the results from three biological replicates each performed with three technical replicates. Statistical significance was assessed via Student’s T-test in all panels except 1A, in which Dunnett’s multiple comparisons post-test was used. *, **, indicate differences that are significantly different at P<0.05 and P< 0.01, respectively, compared to the WT. ns; non-significant.

To verify that the loss of the biofilm that we observed at the 48h assay was dependent on the absence of the *yfiN* gene, we genetically complemented the Δ*yfiN* mutant with a plasmid generously provided by Thomas Wood (32). After induction with arabinose (0.2% w/v) to induce expression of this gene from the plasmid, we found that biofilms of WT carrying the pMQ70 empty vector (EV), the Δ*yfiN* mutant carrying the EV, and Δ*yfiN* mutant complemented with pMQ70-*yfiN* to be comparable at 12h (**Fig. 1A**, left panel). The biofilm defect at 48h observed in the Δ*yfiN* mutant carrying the EV can be largely rescued for the Δ*yfiN* mutant complemented with pMQ70-*yfiN* (**Fig. 1A**, right panel).

**Table S1.**
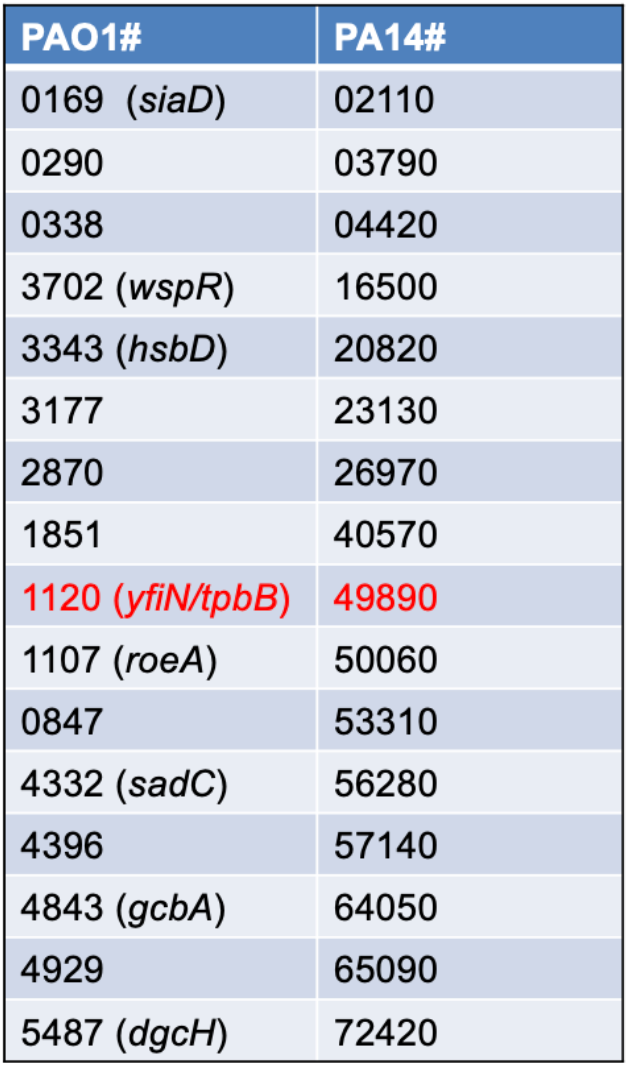
Diguanylate cyclase mutants screened for defects in biofilm maintenance. Previous work by Ha et al. (31) created in-frame, unmarked deletions of all putative diguanylate cyclases that were identified through SMART analyses of proteins encoded in the *P. aeruginosa* genome (31). Here, these mutants were grown statically in 96-well plates for 12h and 48h in M63 biofilm medium. Of all mutants screened, only the Δ*yfiN* mutant (red) exhibited both comparable biofilm biomass to the WT at 12h, yet was unable to maintain a biofilm at 48h.

We next wanted to understand if the observed biofilm defect of the Δ*yfiN* mutant at 48h was elicited by starvation or other consequences of prolonged batch culture growth. To test this idea, we repeated the 48h static biofilm assay but periodically (every 12h) removed the spent medium, washed the biofilm and added fresh medium. Through this medium replacement assay, we observed that the biofilm defect of the Δ*yfiN* mutant was rescued at 48h (**Fig. 1B**, right panel), with the level of biofilm staining comparable to the WT (**Fig. 1B**, left panel).

Considering the observed biofilm maintenance defect of the *yfiN* null mutant and the previous reports indicating that YfiR activity is regulated by oxidative stress (35), we hypothesized that the biofilm maintenance defect manifested by the Δ*yfiN* mutant may be due to an altered response to peroxide stress. To test this hypothesis, we compared the amount of biofilm retained by WT or the Δ*yfiN* mutant after 24h with a single medium replacement at 12h, to biofilms of these same strains exposed to 50 mM H_2_O_2_ added to the replacement medium. After 24h with control replacement medium, biofilms of WT and the Δ*yfiN* mutant were not significantly different (**Fig. 1C**, left panel). However, when replacement medium contained 50 mM H_2_O_2_ we observed that biofilms of the Δ*yfiN* mutant exhibited a significantly lower biofilm biomass compared to the WT (**Fig. 1C**, right panel). Collectively, these results suggest that YfiN may be important for biofilm maintenance and that *P. aeruginosa* biofilms lacking this DGC exhibited reduced biofilm biomass when exposed to peroxide.

### Dispersion and cell death contribute to the Δ*yfiN* mutant biofilm maintenance defect

The above results provide evidence that YfiN impacts biofilm maintenance in the face of peroxide stress. In order to determine biomass and cell viability before and after treatment, static biofilms were grown in glass-well dishes containing the same medium used in the 96-well dish assay to directly image the biofilms. The biofilm biomass was determined by measuring the fluorescence of the WT and the Δ*yfiN* mutant strain carrying the multicopy plasmid pSMC21, which constitutively expresses GFP (ex500/em513). To visualize dead cells, the biofilms were washed with the same culture medium containing propidium iodide (PI) immediately prior to imaging. Fluorescence intensity was used to quantify the biofilm biomass of each strain (green fluorescence), and the ratio of PI to GFP intensity was used as a metric to measure normalized cell death within the biofilms.

Here, the Δ*yfiN* mutant exhibits a mild, but in this case significant, defect in the biofilm biomass at 12h compared to the WT in this glass dish assay (**Fig. 2A, E-F)**. These data suggest that both strains can establish a biofilm. Consistent with the 96-well dish assay (**Fig. 1A**), the Δ*yfiN* mutant shows significantly less biofilm biomass at 24h compared to the WT (**Fig 2B, G, H**). Thus, we can use the glass bottom dishes to visualize the biofilms and largely replicate the findings observed in the 96-well dish assay.

**Figure 2.**
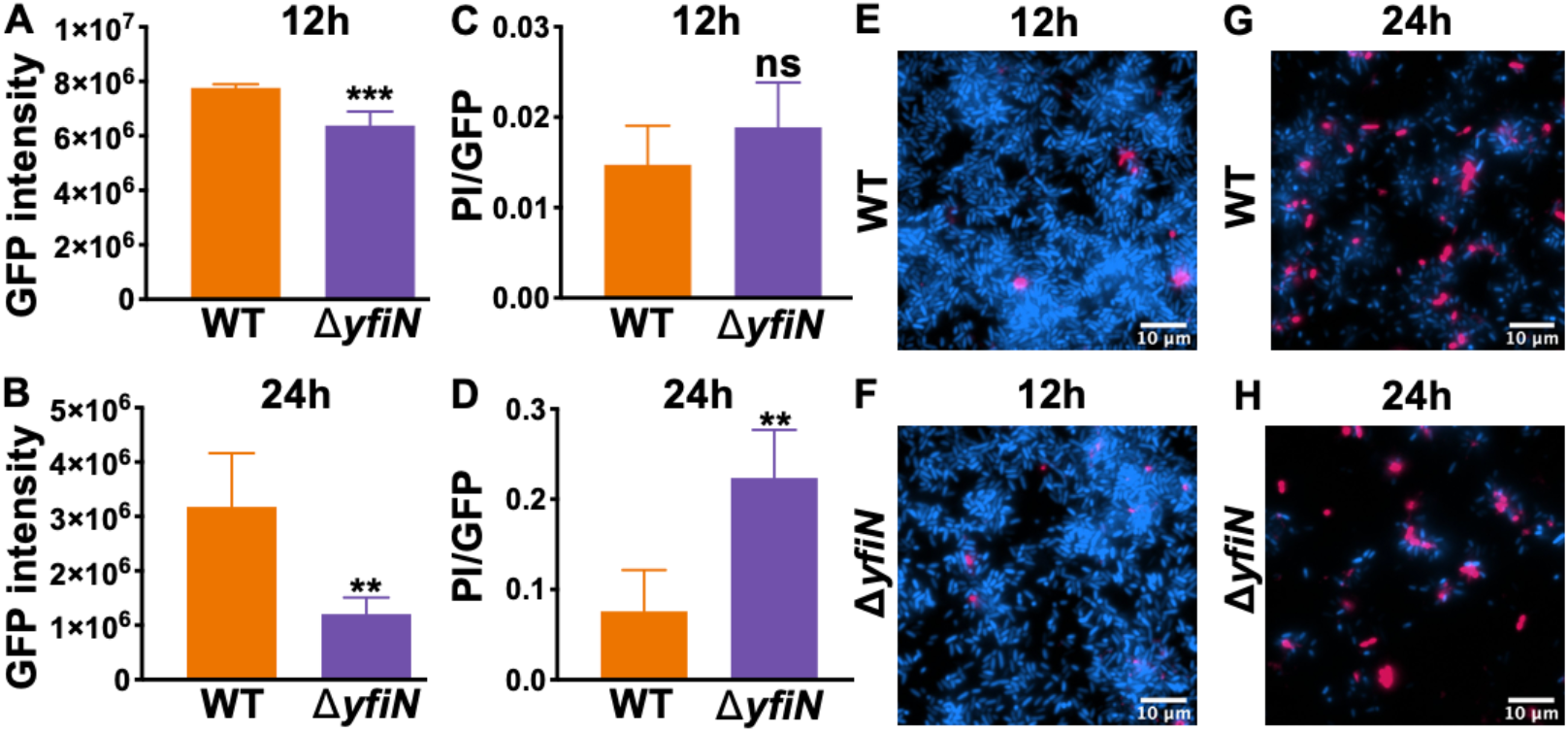
Cell death likely contributes to the Δ*yfiN* mutant biofilm maintenance defect. Biofilm biomass and cell death was assessed microscopically in glass-bottom dishes with WT and the Δ*yfiN* mutant. Cell biomass was determined using fluorescence mediated by the plasmid pSMC21, which encodes a constitutively-expressed GFP. Cell viability was determined as a ratio of propidium iodide (PI)-stained (dead) cells to GFP-mediated green fluorescence (live cells). The PI was added to the dishes immediately prior to visualizing, and the cells were washed to remove excess dye. A-B) WT and the Δ*yfiN* mutant biofilm biomass was quantified by measuring GFP intensity at 12h (A) and 24h (B). We observed that the biofilm maintenance defect of the Δ*yfiN* mutant occurred as early as 24h in the glass-bottom plates assay and consequently used this point for the analysis here. C-D) Cell death was measured as a ratio of PI staining to GFP intensity at 12h (C) and 24h (D). For all panels, error bars represent standard deviations of the results from three biological replicates each performed with 6 technical replicates. Statistical significance was assessed via a Student’s T-test. **, ***, indicate differences that are significantly different at P< 0.01 and P<0.001 respectively, compared to the WT. ns, non-significant. (E-H) Representative images showing combined GFP (blue) and PI staining (pink) of WT biofilm at 12h (E) and 24h (G), and the Δ*yfiN* mutant biofilm at 12h (F) and 24h (H).

When we assessed cell viability via the ratio of PI-stained dead cells to GFP-labeled live cells, we observed no difference between the WT and Δ*yfiN* mutant at 12h (**Fig. 2C**). While we did detect dead (PI-stained) cells for the WT strain at 24h, the extent of cell killing as assessed by the PI/GFP ratio was significantly higher for the Δ*yfiN* mutant (**Fig. 2D)**. Representative images used to generate the quantitative data in panels A-D are shown in **Fig. 2** panels E-H. Together, these data suggest that the loss of the biofilm in the *yfiN* mutant at 24h is due, at least in part, to increased cell death in this mutant.

### The Δ*yfiN* mutant biofilm exhibits increased susceptibility to peroxide stress under flow

The static biofilm data presented thus far are consistent with the hypothesis that YfiN contributes to biofilm maintenance and that the loss of this DGC enhances cell death as these communities age and/or encounter oxidative stress in the form of peroxide. We next wanted to determine the impact of H_2_O_2_ specifically on the Δ*yfiN* mutant biofilm under flow conditions, which helps mitigate the presence of potential confounding factors, such as accumulating metabolites in the extracellular milieu. To address this question, we pursued a microfluidics-based approach in which WT and the Δ*yfiN* mutant with constitutive expression of GFP (using pSMC21, described above) were grown as a biofilm under flow in biofilm medium, a buffered minimal medium containing 1.0 mM K_2_HPO_4_, 0.6 mM MgSO_4_ and 0.4% arginine, pH=7.5. This medium is similar to the M63-based medium used in the static assay but has been optimized for use in flow studies.

First, to determine if the loss of YfiN adversely impacted the ability of this mutant to form a biofilm under standard flow conditions (i.e., no H_2_O_2_ added), we compared GFP intensity with that of WT over a 12h time course kinetic assay. The WT and Δ*yfiN* mutant biofilms were comparable for the first 10h of this assay, after which WT biofilm was modestly but significantly higher than that of Δ*yfiN* mutant in three of the last five time points assessed after 10h (**Fig. S1**).

**Figure S1.**
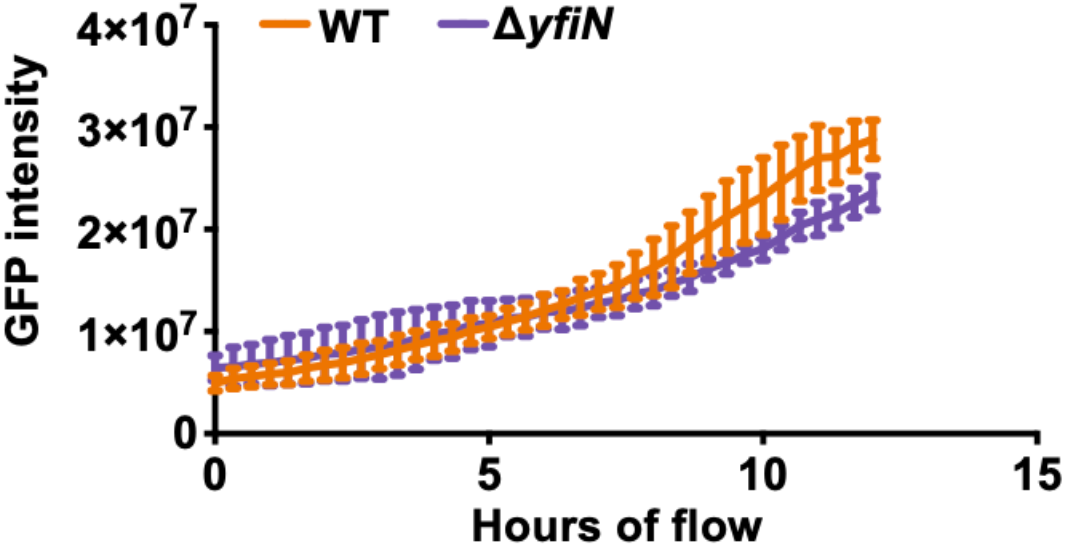
Initial biofilm formation by WT and the Δ*yfiN* mutant under flow. The WT strain and the Δ*yfiN* mutant carrying pSMC21, a plasmid which constitutively expresses GFP, were inoculated into microfluidic chambers and allowed 1h to attach prior to beginning flow of biofilm medium. Flow was resumed, then GFP fluorescence was measured every 20 minutes for 12h. Error bars represent standard deviations of the results from three biological replicates each performed with three technical replicates. Statistical significance was assessed via Sidak’s multiple comparisons post-test. The WT formed significantly more biofilm at 10.7, 11 and 11.7h.

We first assessed whether the Δ*yfiN* mutant was more susceptible to starvation than the WT, as was reported for strains carrying mutations in the *rmcA* or *morA* genes (30); we observed no such phenotype in the microfluidic devices (not shown). To determine whether the Δ*yfiN* mutant was more susceptible to H_2_O_2_ under flow compared to the WT, both strains were grown for 14h as a biofilm, after which influent biofilm medium was replaced with medium containing 10 mM H_2_O_2_, as well as PI to stain dead cells, and the biofilms imaged over an additional 12h time course. We then assessed biofilm biomass, cell viability and cell dispersion after this 12h exposure to peroxide.

Using GFP signal intensity as a measure of biofilm biomass, we found that prior to the addition of H_2_O_2_, there was no significant difference in the biofilm biomass of the WT versus the Δ*yfiN* mutant over the first 10h of the experiment (**Fig. S1**), indicating that both strains could form a robust biofilm. We did note a difference in the biofilm biomass between these strains under flow after 12h, but this reduction was modest and not significant at every time point (see **Fig. S1**). Monitoring the biofilm biomass showed that the addition of H_2_O_2_ resulted in a significant decrease in Δ*yfiN* biofilm biomass 6h after treatment compared to the WT, which continued to increase in biomass as judged by GFP signal intensity (**Fig. 3A, D**). Our analysis shows that H_2_O_2_ treatment adversely impacted the viability of Δ*yfiN* mutant; this mutant exhibited a significant increase in its PI/GFP ratio compared to WT starting 3h after H_2_O_2_ treatment and at each subsequent time point (**Fig. 3B, D**).

**Figure 3.**
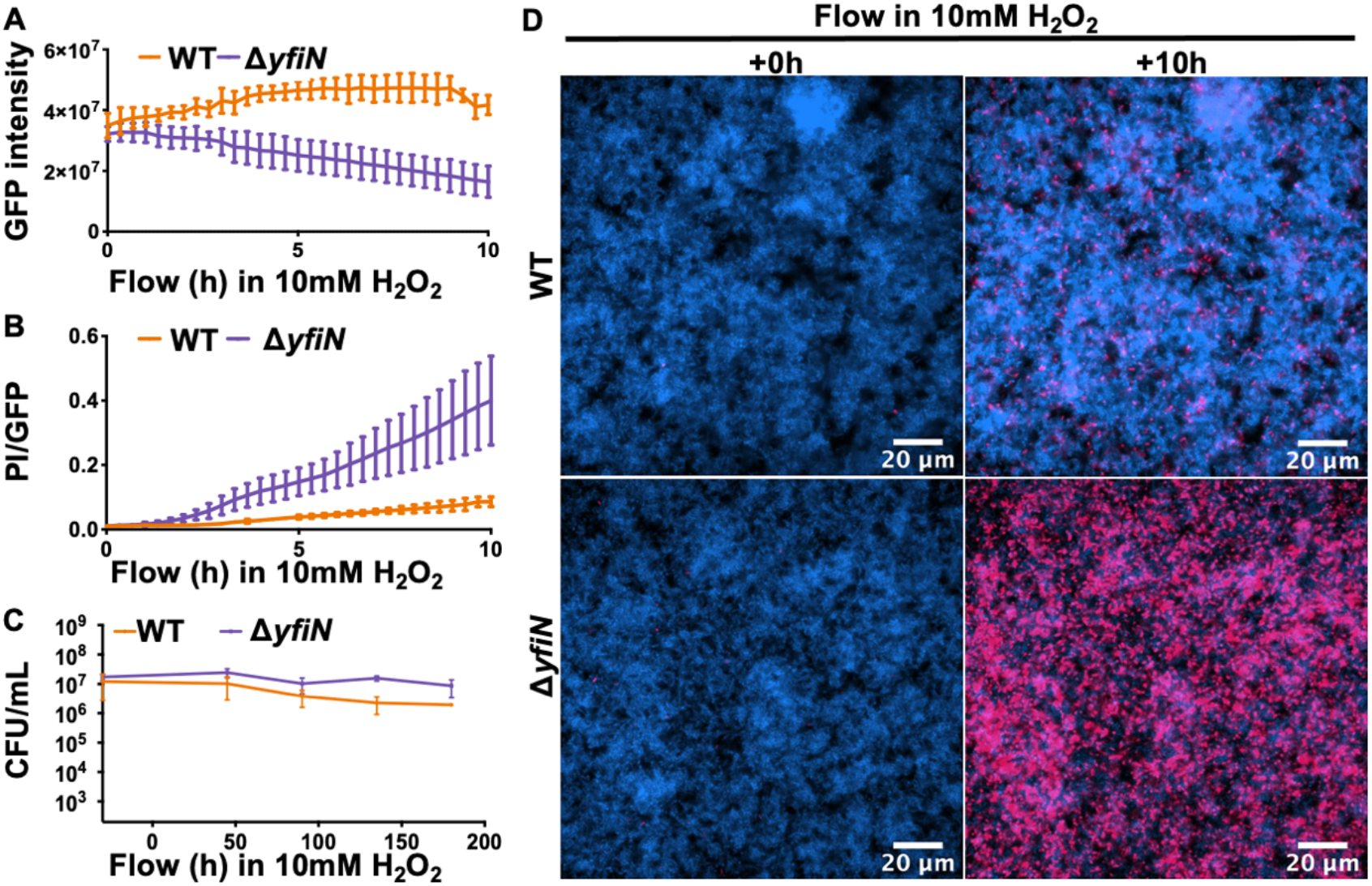
The Δ*yfiN mutant* biofilm exhibits increased cell death and dispersal in response to peroxide treatment. A) Biofilm formation by the WT and the Δ*yfiN* mutant was quantified through fluorescent microscopy and GFP signal intensity measured for 10h after the addition of H_2_O_2_. B) Cell death was measured as a ratio of PI to GFP over the same time course as panel A. C) WT and the Δ*yfiN* mutant carrying plasmid pSMC21 encoding GFP were inoculated into a microfluidics device and grown in biofilm medium. After 14h of growth, medium was replaced with biofilm medium containing 10 mM H_2_O_2_ and PI. Bacterial CFUs/mL were measured in the effluent 30 min prior to and up to 180 min after being exposed to H_2_O_2_. Error bars represent standard deviations of the results from three biological replicates each performed with three technical replicates. Statistical significance was assessed via Sidak’s multiple comparisons post-test. Compared to WT, results from the Δ*yfiN* mutant were significantly different (P< 0.05) from WT after 6h in Figure 3A, 3h in Figure 3B, and at 45 min and 120 min in Figure 3C. D) Representative images from microfluidics studies used to generate the quantitative data presented in panels A and B. Viable cells (GFP) are colored blue while PI-stained cells are colored magenta.

To determine if H_2_O_2_ exposure caused cells to disperse from the biofilm, colony forming units (CFUs) were measured in the effluent 30 min prior to and for 3h after the H_2_O_2_ -containing medium switch. Prior to exposure to H_2_O_2,_ cell dispersal between WT and mutant Δ*yfiN* were not significantly different. However, the switch to medium containing 10mM H_2_O_2_ resulted in modest increases of Δ*yfiN* mutant cells in the effluent, and this difference was significant at 45 and 120 min post-stress exposure (**Fig. 3C**). Collectively, the use of static and microfluidics assays showed that the loss of YfiN function resulted in increased cell death, as well as a modest increase in dispersal of the biofilm cells compared to the WT.

To test if the enhanced H_2_O_2_ sensitivity exhibited by the Δ*yfiN* mutant was specific to the biofilm lifestyle, we washed and resuspended overnight-grown planktonic cultures of the WT and the Δ*yfiN* mutant and challenged these strains with 10mM H_2_O_2_ for 4 hours. There was no significant difference between the viability of the WT and the Δ*yfiN* mutant (**Fig. S2**), indicating the sensitivity to H_2_O_2_ was biofilm-specific.

**Figure S2.**
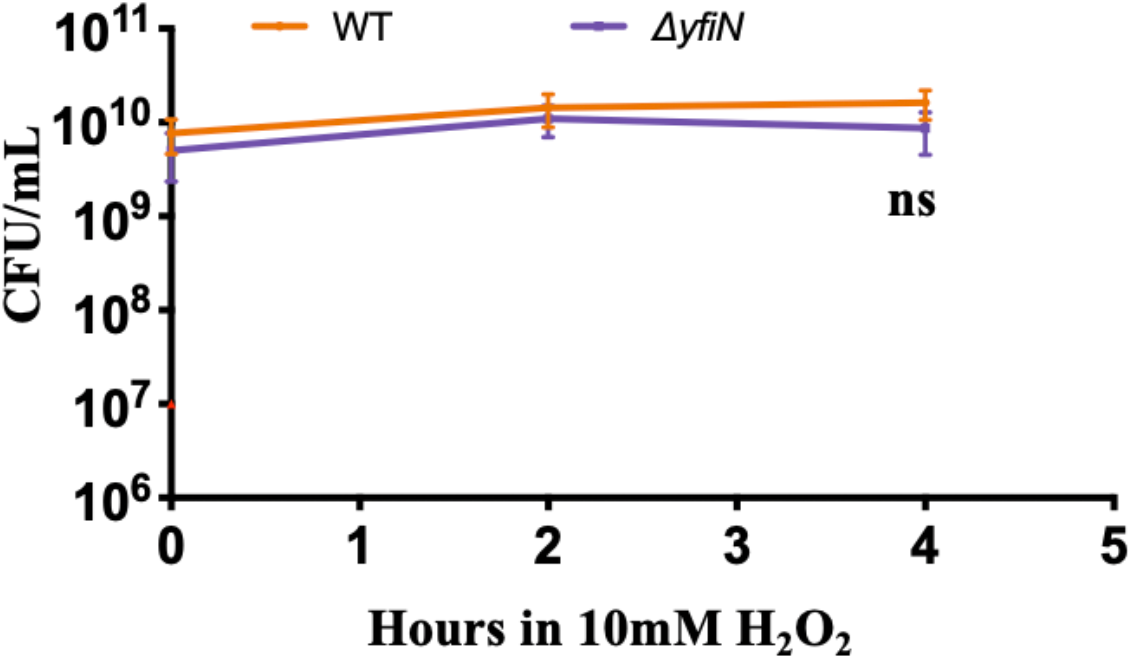
The Δ*yfiN* mutant does not exhibit a significant increase in H_2_O_2_ susceptibility during planktonic culture. The WT strain and the Δ*yfiN* mutant were inoculated into biofilm medium, grown for 24h, washed, re-suspended in fresh medium with and without 10mM H_2_O_2_ to a normalized OD_600_ of 0.5. CFU/mL were measured prior to and 2-4h after the addition of H_2_O_2_. Error bars represent standard deviation of the results from three biological replicates each performed with three technical replicates. Statistical significance was assessed with a Sidak’s multiple comparisons post-test; ns, indicates a non-significant.

### Loss of YfiN impacts c-di-GMP levels and motility in response to peroxide stress

YfiN contains a DGC domain that has been shown to mediate the synthesis of c-di-GMP (32, 33). We next assessed whether exposure to H_2_O_2_ impacted c-di-GMP levels. To measure c-di-GMP level, we used a P_*cdrA*_-*gfp* promoter fusion on a plasmid to assess the transcriptional activity of *cdrA*, a gene that is positively regulated by c-di-GMP. We normalized the GFP signal to biomass using mKO (ex548/em559) inserted as a double copy at the *att* site on the chromosome and expressed constitutively. Bacteria were inoculated into glass-bottom dishes, grown as a monolayer of surface-attached cells, exposed to 1mM H_2_O_2_ (a concentration of peroxide that is well below the toxic levels analyzed above) and then imaged over the next 8h. Consistent with previous findings, the loss of YfiN reduced c-di-GMP levels, evident in the lower GFP signal (pink in images) for the Δ*yfiN* mutant compared to WT (**Fig 4A**, top left and bottom left, respectively, and **Fig. 4B**, time = 0). After exposure to peroxide, we observed an initial drop in GFP intensity by both WT and the Δ*yfiN* mutant (**Fig. 4B**), indicative of less c-di-GMP in response to hydrogen peroxide. Interestingly, while WT exhibited a recovery in c-di-GMP after ∼6h, this recovery was not observed in the Δ*yfiN* mutant, for which the GFP signal remained low.

**Figure 4.**
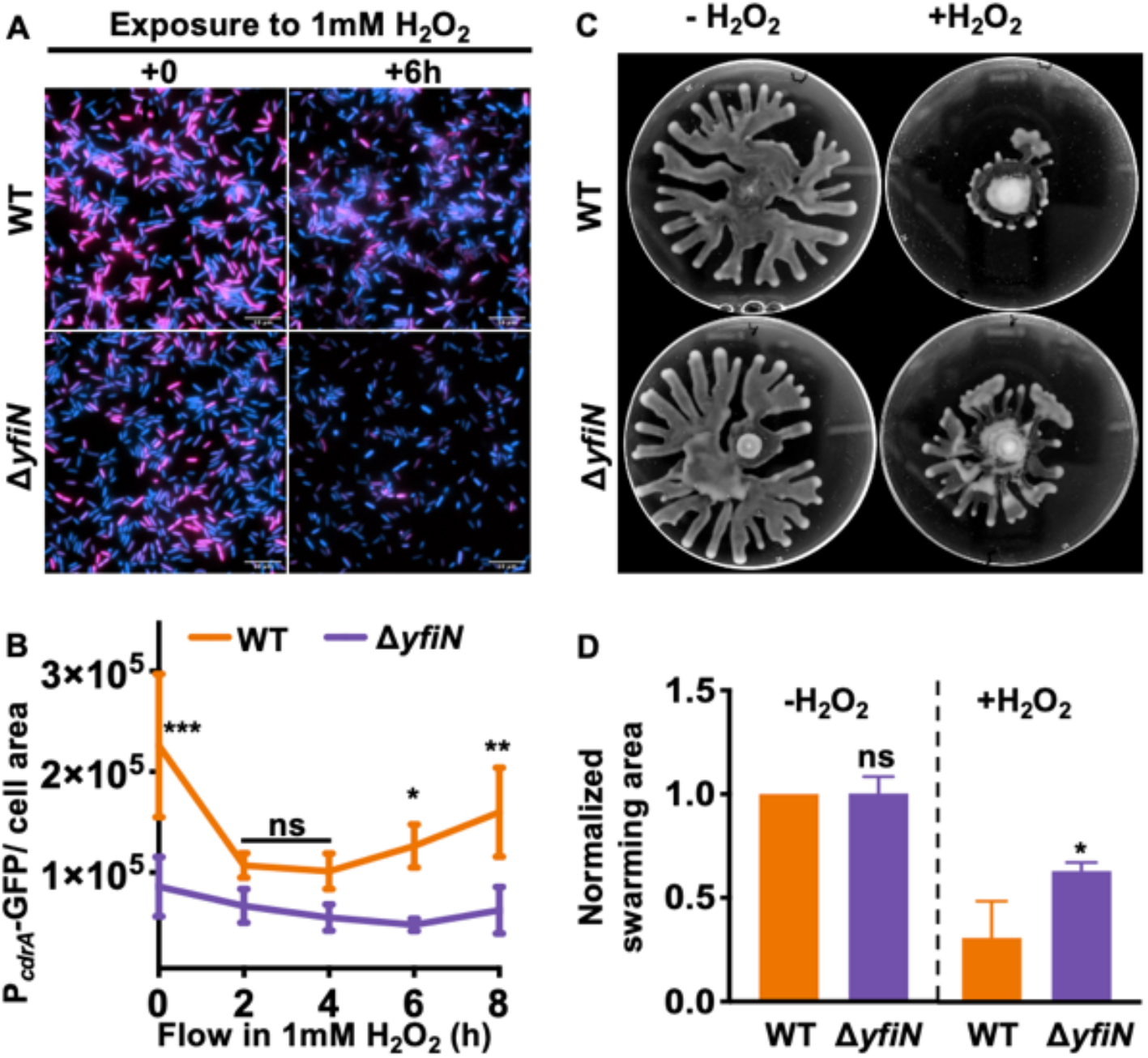
Loss of YfiN impacts c-di-GMP levels and motility in response to peroxide stress. A) The WT strain and the Δ*yfiN* mutant carrying the P_*cdrA*_-*gfp* fusion expressed from a multicopy plasmid were grown in glass bottom 8-well dishes containing M63 medium for 3h prior to the addition 1 mM H_2_O_2_, then imaged over 8h with a representative image from the 0h and 6h time points shown. The pink color shows GFP expression, and the blue color is contributed by the constitutively expressed mKO fluorescent protein. (B) Quantification of GFP signal intensity for the strains described in panel A. Fluorescent microscopy was used to determine GFP signal intensity as a measure of c-di-GMP production and normalized to the surface area of cells that constitutively expressed mKO fluorescent protein. Statistical significance was assessed with a Sidak’s multiple comparisons post-test. C) Representative images of the indicated strains grown on swarm assay plates 16h after inoculation, with and without 5 mM H_2_O_2_. D) Quantification of plates described in panel C; all values are plotted after normalization to the WT not treated with H_2_O_2_, which was set to a value of 1. For panels B and D, error bars represent standard deviations of the results from three biological replicates each performed with three technical replicates. *, **, *** indicate differences that are significantly different at P<0.05, P< 0.01, and P< 0.0001 respectively, compared to the WT. ns, non-significant.

Two key mechanisms whereby c-di-GMP regulates biofilm formation and maintenance are through (1) mediating increased production of extracellular polysaccharide (EPS) and/or (2) limiting motility. Previous findings from our lab did not observe a decrease in EPS production in the Δ*yfiN* mutant (31). To determine if the reduced biofilm phenotype in response to H_2_O_2_ observed for the Δ*yfiN* mutant was caused, at least in part, by increased motility in response to H_2_O_2_, we first measured distance traveled by single cells imaged on glass well plates. Prior to the addition of 5mM H_2_O_2_ the motility of WT and the Δ*yfiN* mutant was comparable (**Fig. S3, Movies S1** and **S2**). Thirty minutes after the addition of H_2_O_2_, the motility of the Δ*yfiN* mutant was significantly higher than the WT (**Fig. S3, Movies S3** and **S4**).

**Figure S3.**
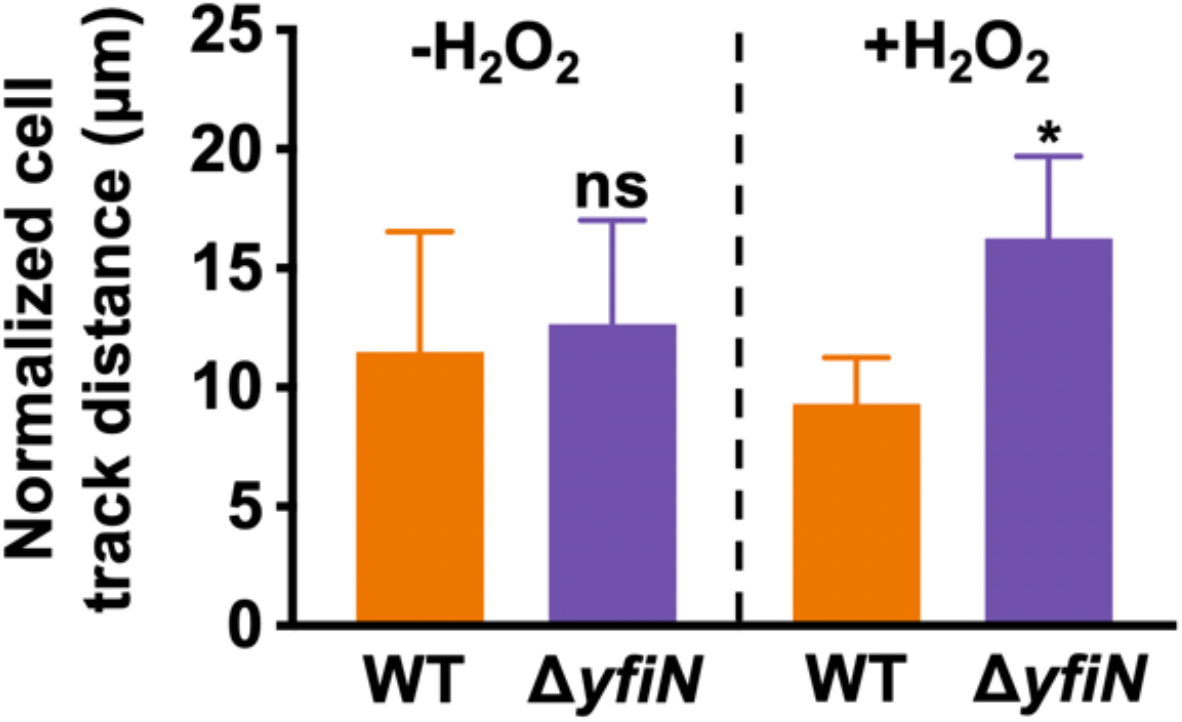
The Δ*yfiN* mutant exhibits significant increased motility in response to H_2_O_2_. The WT strain and the Δ*yfiN* mutant were inoculated into glass well dishes for 1h. Motility was assessed over a 5 minute time course using brightfield microscopy prior to and 30 minutes after the addition of 5mM H_2_O_2_. The TrackMate program was used to measure the tracks that each bacterium traveled over time, and the sum distance of all cell tracks was normalized to the number of bacteria at the start of imaging. Statistical significance was assessed via a Student’s T-test. * indicates differences that are significantly different at P<0.05, compared to the WT. ns, non-significant.

**Movie S1.**
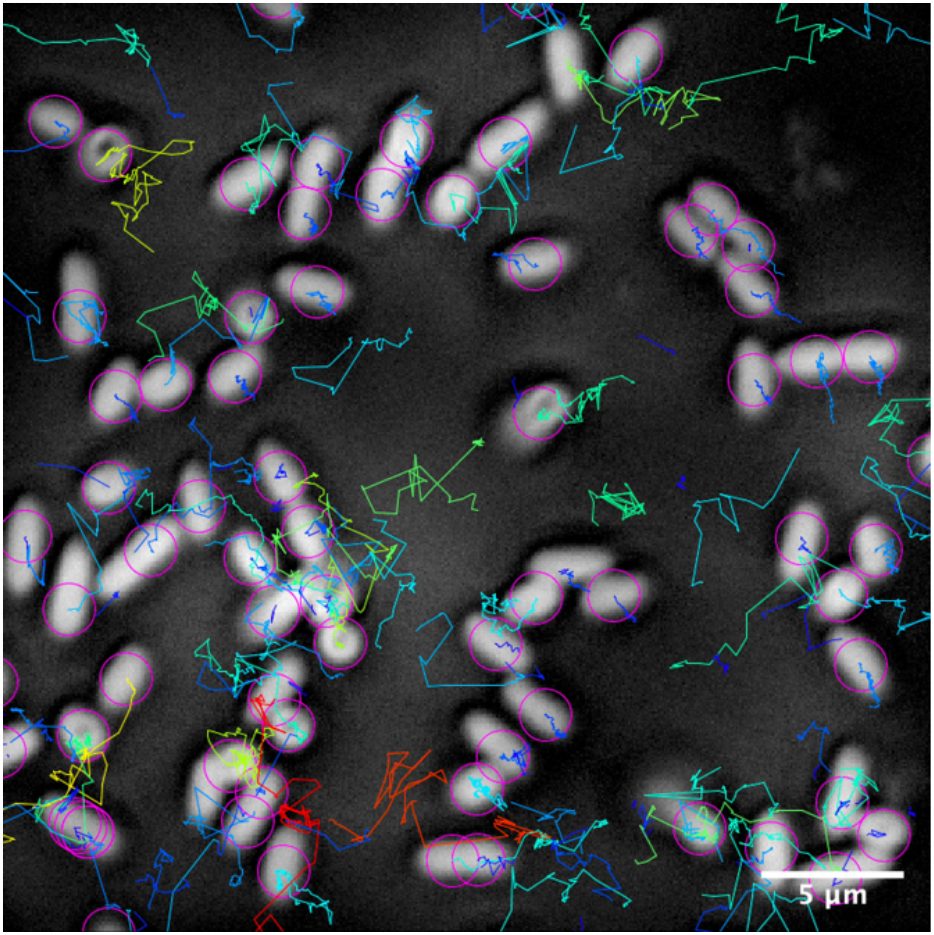
Motility of WT prior to the addition of H_2_O_2._ WT was inoculated into a glass well dish. Surface-attached bacteria were imaged one hour after inoculation with brightfield microscopy over a 5 minute time-course. The TrackMate program was used to identify bacteria (purple circles) and the distance of the tracks that bacteria traveled over time was measured and bacterial tracked distance depicted as a color-coded spectrum (blue<10µm; red>30µm).

**Movie S2.**
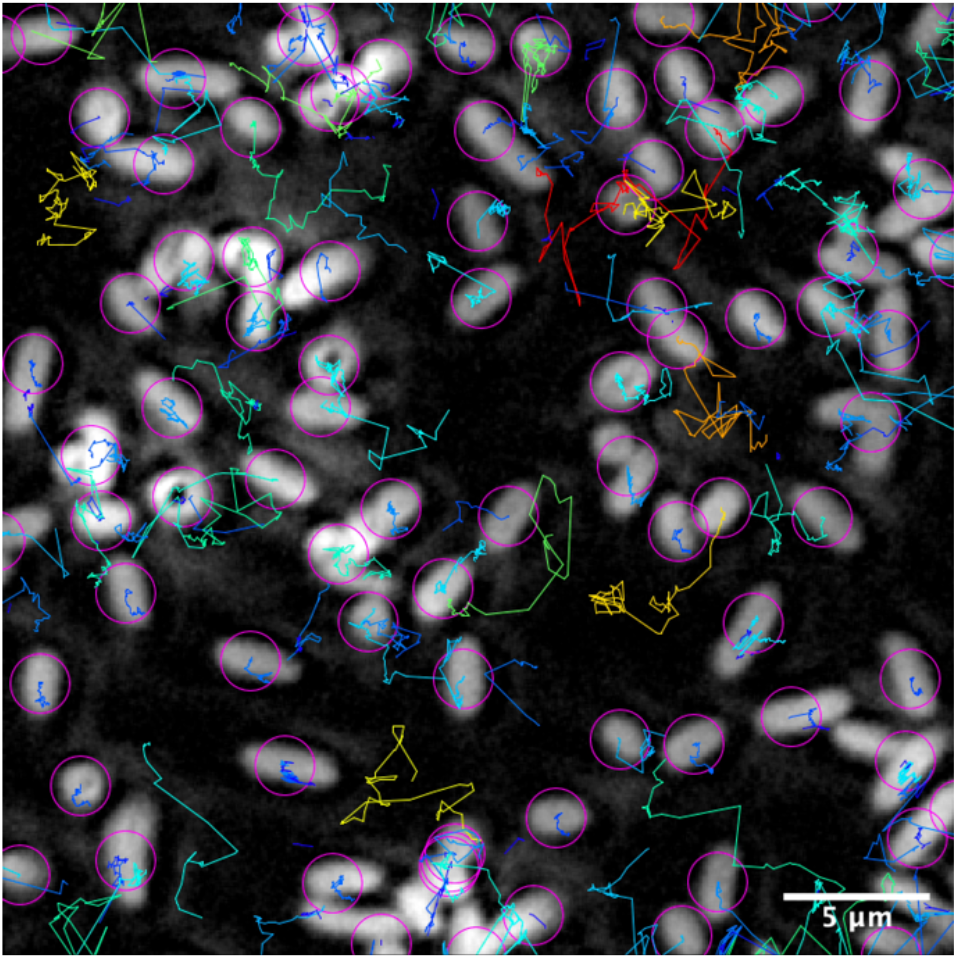
Motility of the Δ*yfiN* mutant prior to the addition of H_2_O_2._ The Δ*yfiN* mutant was inoculated into a glass well dish and surface-attached bacteria were imaged 1h after inoculation, as described in Movie S1. The distance of the tracks that bacteria traveled over time was measured and bacterial tracked distance depicted as a color-coded spectrum (blue<10µm; red>30µm).

**Movie S3.**
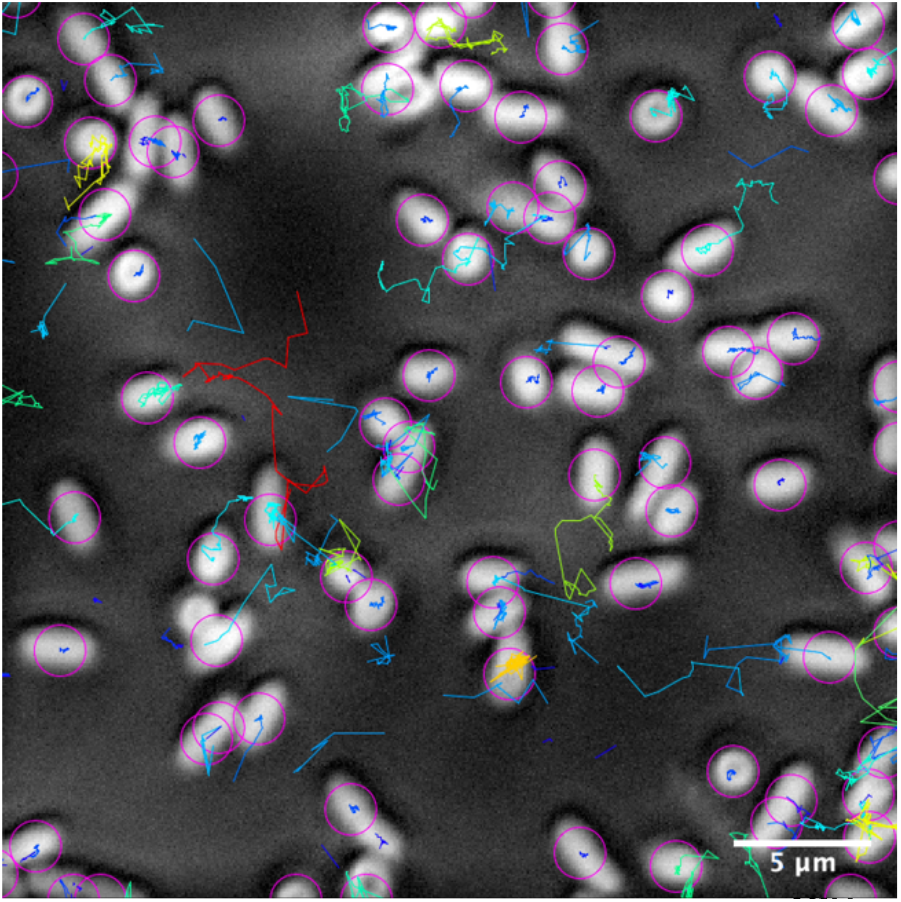
Motility of WT after the addition of H_2_O_2._ WT was inoculated into a glass well dish and allowed to attach to the surface for 1h. Bacteria were subsequently exposed to 5mM H_2_O_2_ and imaged after 30 minutes for a 5 minute time-course using brightfield microscopy. The distance of the tracks that bacteria traveled over time was measured and bacterial tracked distance depicted as a color-coded spectrum (blue<10µm; red>30µm).

**Movie S4.**
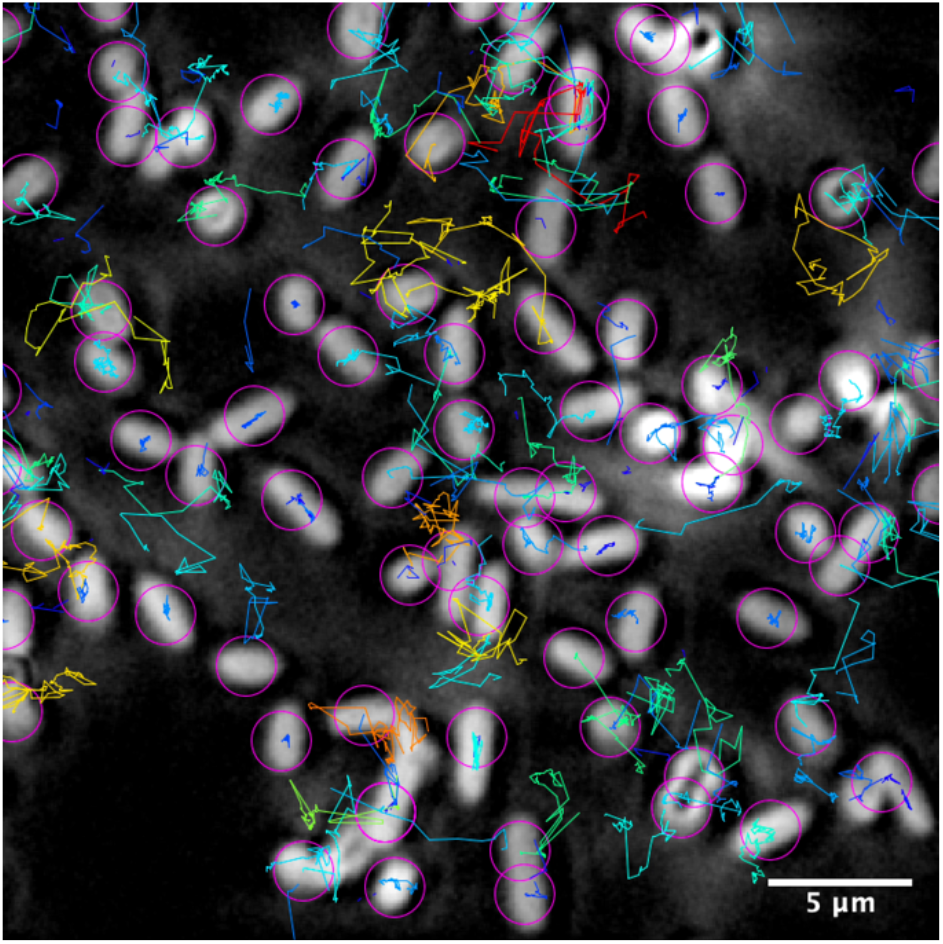
Motility of the Δ*yfiN* mutant after the addition of H_2_O_2._ The Δ*yfiN* mutant was inoculated into a glass well dish and surface-attached bacteria were imaged 1h after inoculation as described in Movie S1. The distance of the tracks that bacteria traveled over time was measured and bacterial tracked distance depicted as a color-coded spectrum (blue<10µm; red>30µm).

We next assayed swarming on 0.5% agar plates in response to +/-5mM H_2_O_2_. Swarming motility by *P. aeruginosa* is a surface-based motility which utilizes the flagellum. In the absence of H_2_O_2_, both strains exhibit comparable swarm motility (**Fig 4C-D**, left panels). While the addition of 5mM H_2_O_2_ resulted in almost complete suppression of swarming motility by the WT, the Δ*yfiN* mutant exposed to H_2_O_2_ still displayed swarming motility, although somewhat reduced from the no treatment control (**Fig 4C-D**, compare right panels to left panels). Collectively, these results suggest that *P. aeruginosa* uses YfiN to modulate c-di-GMP levels and suppress motility when exposed to peroxide stress.

### Likely YfiBNR homologs are found in other bacterial genera

Known YfiN homologs are present in other biofilm-forming, Gram-negative bacteria like *Pseudomonas fluorescens, Escherichia coli*, and *Yersinia pestis* (36–39). Given the above evidence that YfiN regulates biofilm maintenance in *P. aeruginosa*, we mined GenBank for YfiBNR homologs in other bacteria, with a particular focus on Gram-negative, biofilm forming microbes. To identify potential homologs, we used the nucleotide sequence of the *P. aeruginosa* YfiBNR operon as a query in an NCBI Nucleotide BLAST search against the RefSeq representative genomes database (40). We aligned the 280 sequences with >75% query cover via Multiple Alignment using Fast Fourier Transform (MAFFT), then mapped sequences into a phylogenetic tree using FastTree 2.1.11 (41, 42).

The resulting phylogenetic tree (summarized in **Fig. 5A**) does not group by taxonomic classification and contains 116 species of bacteria in eight genera: 83.2%, 7.9%, and 3.9% of the hits were *Pseudomonas, Janithobacterium*, and *Serratia*, respectively. *Cupriavidus, Rugamonas, Inquilinus, Bordetella*, and *Achromobacter* each made up 0.4-1.4% of the remaining sequences. The resulting phylogenetic tree identified a cluster (**Fig. 5B)** with high bootstrap support that formed around the query sequence. This cluster contains *Pseudomonas* and *Serratia*, which are genera of different orders. Notably, the cluster includes a sequence from *Serratia marcescens*, a known biofilm-forming, opportunistic human pathogen that infects the respiratory tract, urinary tract and eyes (43–45). These YfiBNR homologs may be similarly involved in biofilm maintenance and response to peroxide stress in their respective species.

**Figure 5:**
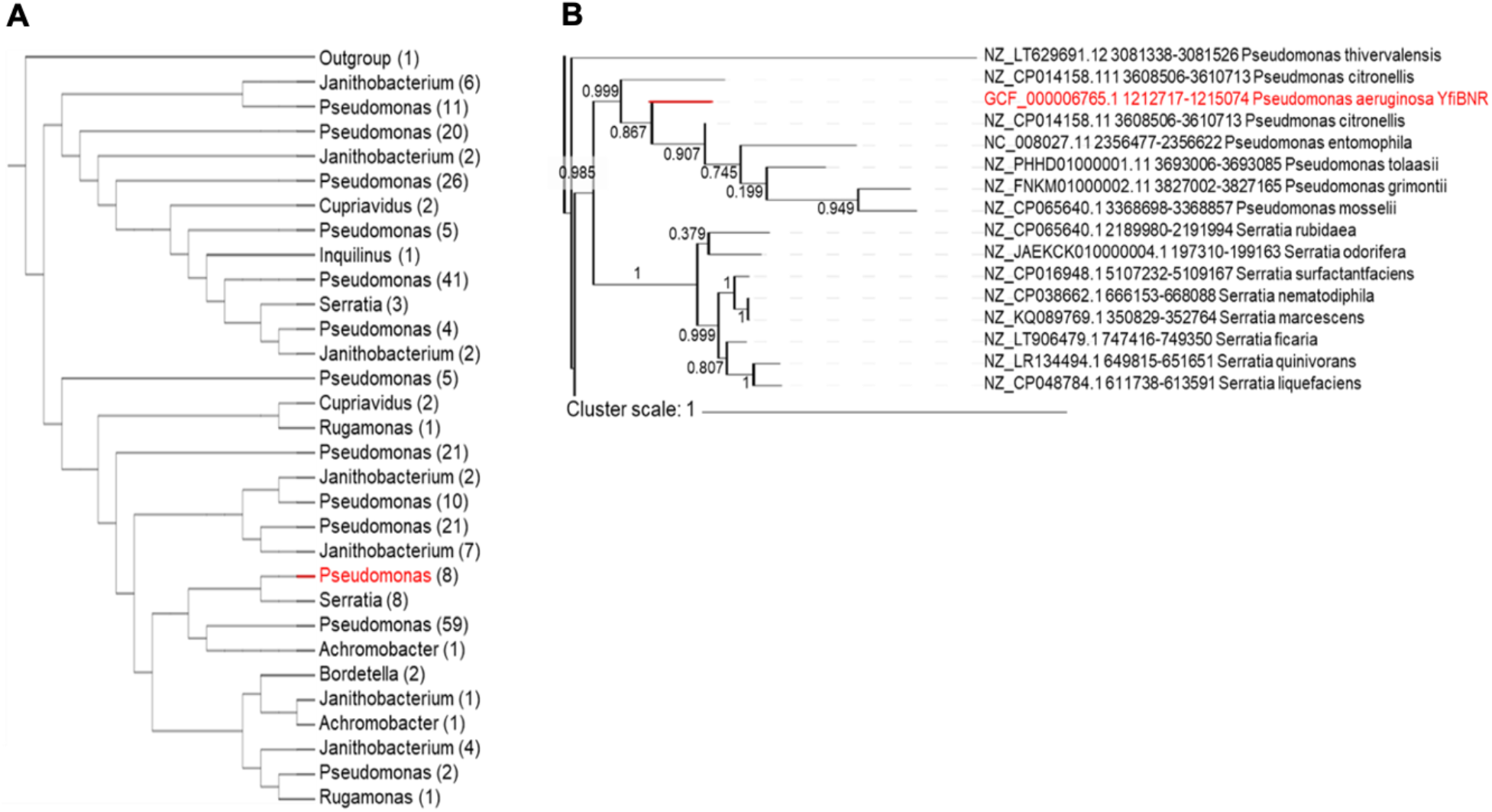
YfiBNR homologs are found in multiple bacterial genera. A) Potential YfiBNR homologs were identified via a Nucleotide BLAST search, aligned with MAFFT, arranged into a phylogenetic tree using FastTree 2.1.11, and visualized using Interactive Tree of Life software (46). Sequences most closely mapped to sequences of the same genus were collapsed to generate the above abbreviated tree. The full tree is shown in **Figure S4**. Each branch is labeled with the number of sequences from that genus present on that branch in the full tree. The branch containing the *P. aeruginosa* YfiBNR query sequence is marked in red. Branch lengths are not mapped on this tree. B) A cluster labeled with high bootstrap values, indicating frequent mapping of the same branches in various iterations of the tree, formed around the YfiBNR query sequence, which is marked in red. Branch lengths, which mark how much genetic divergence exists between sequences, are to scale. The sequences mapped most closely to YfiBNR were from multiple other *Pseudomonas* and *Serratia* strains, suggesting potential homology across species and genera.

**Figure S4.**
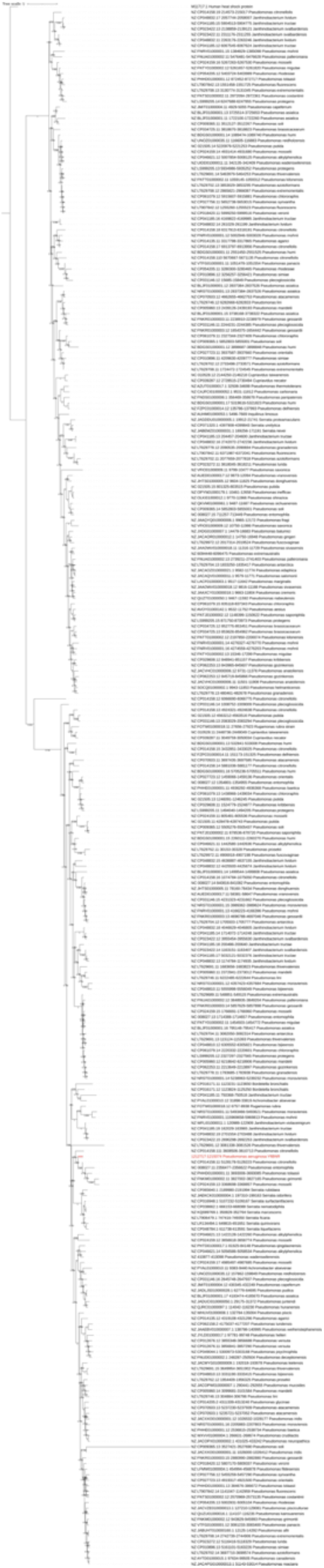
Full phylogenetic tree of potential YfiBNR homologs. The full phylogenetic tree of potential YfiBNR homologs was produced from 280 aligned sequences from 116 species in eight genera.

## Discussion

YfiN homologs are found among various bacterial genera, including *Pseudomonas, Escherichia, Salmonella* and *Serratia*, and this protein has been shown to play a role in regulating intracellular c-di-GMP in *P. aeruginosa* here and in previous studies (32, 47). The contributions of YfiN have been largely studied in the context of motility and biofilm formation, whereby a loss of YfiN leads to less c-di-GMP and reduced biofilm formation (32, 48), consistent with our results in *P. aeruginosa*. Increased motility and the resulting dispersal of the biofilm that we observe here has been indirectly supported through the study of upstream regulators. For instance, deletion of YfiR, an upstream inhibitor of YfiN, has been shown to repress motility in *P. aeruginosa* PAO1 (33). A release of inhibition by YfiR leads to activation of the c-di-GMP-producer YfiN, thus reducing motility. Furthermore, previous biochemical studies have shown that oxidized glutathione disrupts disulfide bond formation and protein stability of YfiR (35), suggesting the critical role of redox stress on the regulation of this pathway.

Here we observe a significant biofilm maintenance defect at 48h, but not 12h, in the Δ*yfiN* mutant. Consistent with the loss of this DGC, we observe a significant decrease in *cdrA* transcription, an indirect read-out of c-di-GMP level, in the Δ*yfiN* mutant compared to WT. We also note increased cell death and a modest increase in dispersion in the Δ*yfiN* mutant compared to WT. Thus, we attribute this loss of biofilm maintenance both to enhanced dispersal and increased cell death. We provide evidence that YfiN modulates c-di-GMP levels, cellular motility and the ability to maintain the biofilm in response to peroxide (**Fig. 6**). At this stage, we have not established the link between the loss of YfiN and the observed cell death and dispersion phenotypes. Low levels of c-di-GMP are associated with increased surface motility of *P. aeruginosa*, with our group and many others contributing to the known mechanisms (12, 18, 49, 50). We presume that c-di-GMP-mediated regulation of motility is through one of the known mechanisms, but this remains to be demonstrated. Why we observe increased cell death in the *yfiN* mutant treated with peroxide is a mystery. Our previous work strongly suggested that loss of RmcA/MorA contributed to cell death during starvation because strains lacking these proteins continued to use resources to make EPS even when starving; the mutants were likely starving themselves to death (30). Here, we suggest two possibilities for the cell death observation. The first is that high c-di-GMP levels promoted by YfiN are necessary to induce protective responses to peroxide. Increased c-di-GMP in *Vibrio cholerae* has been found to induce matrix production and enhance catalase activity (51, 52). In further support of this view in *P. aeruginosa*, the DGC PA3177 has been found to synthesize c-di-GMP in response to oxidative hypochlorite (53) and to mediate H_2_O_2_ and antibiotic tolerance (54). Furthermore, sublethal concentrations of H_2_O_2_ have been shown to select for mutations in the gene encoding for WspF, which represses the DGC WspR (55). The second possibility is that exposure to peroxide in the absence of YfiN results in the biofilm bacteria returning to a less tolerant, planktonic lifestyle. This second hypothesis is supported by the observation that YfiN mutants are more motile and more readily dispersed from the biofilm subsequent to peroxide exposure.

**Figure 6:**
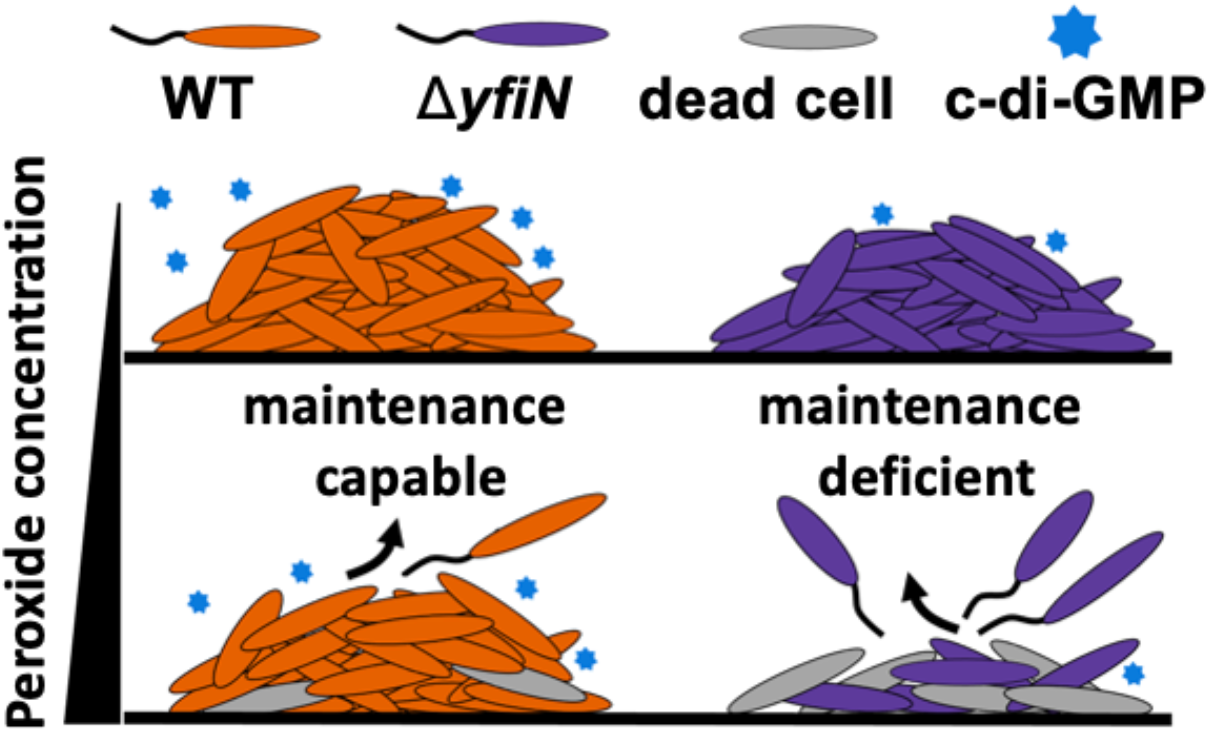
Model *for P. aeruginosa* YfiN’s contribution to biofilm maintenance in response to peroxide exposure. YfiN is not a major contributor to early biofilm formation in our growth conditions (top panel). In a mature WT biofilm (bottom panel), YfiN is needed to regulate maintenance through the modulation of c-di-GMP and motility in response to peroxide. The loss of YfiN results in biofilms that are maintenance-deficient and exhibit increased cell death and dispersal upon exposure to exogenous peroxide, and perhaps in response to endogenous redox active molecules.

Interestingly, we found that loss of biofilm biomass in later-stage biofilms in strains lacking YfiN function in both the absence and presence of H_2_O_2_, and furthermore, the loss of biofilm biomass by the Δ*yfiN* mutant in the absence of H_2_O_2_ could be rescued by replenishing biofilms with fresh medium. These data suggest that YfiN may be important for adapting to oxidative stressors generally present in the environmental milieu, including self-produced redox active compounds produced by *P. aeruginosa*. Phenazines, small redox active molecules that can act as alternate electron acceptors, may be one such molecule. Interestingly, phenazines have been linked to EPS production via the DGC RmcA (56, 57) and we have shown that RmcA is critical for biofilm maintenance in response to starvation (30), thus there are multiple lines of evidence linking phenazines to the control of biofilm biology. Further work is needed to understand if the ability for YfiN to regulate biofilm maintenance is specific to peroxide exposure or if YfiN is more broadly required across a range of redox active agents, including phenazines and immune system-generated molecules with similar chemistry (58–62).

Overall, our data support the model we proposed previously (30), whereby *P. aeruginosa* actively maintains a mature biofilm even in the face of environmental stressors. That is, *P. aeruginosa* appears to utilize at least three different c-di-GMP metabolizing enzymes, RmcA, MorA and YfiN, to actively maintain the biofilm in response to environmental perturbations, including starvation or peroxide exposure. Based on work here and our previous study of RmcA and MorA, the loss of the ability or modulate c-di-GMP in a mature biofilm in the face of environmental stressors has dire consequences for the cell, that is, death. If one also considers that published work showing that *P. aeruginosa* has pathways for the generation of “maintenance energy” in the context of mature biofilms (63–65), it appears that this microbe has a robust set of mechanisms to sense and respond to environmental inputs when in a biofilm, and then regulate its response(s) appropriately. Such biofilm maintenance pathways might be key to preventing biofilm bacteria from “over-reacting” to transient stresses; thus mediating persistence through the perturbation without transitioning to a planktonic state. That is, perhaps biofilms have a type of “fight” or “flight” response regulated by environmental inputs and c-di-GMP levels. We speculate that such “biofilm maintenance” pathways are likely present in other microbes, and furthermore, at least a subset of c-di-GMP-related enzymes for which there is no assigned function play a role in such pathways.

A number of microbes have been shown to have the *yfi* operon, which encodes the tri-partite regulatory system of YfiN, YfiR and YfiB. The current model suggests that YfiR is localized to the periplasm, where it directly inhibits YfiN. This inhibition is relieved in the presence of oxidative stress, which leads to misfolding of YfiR. Additionally, YfiB potentially sequesters YfiR following cell envelope stress, thus relieving inhibition of YfiN (47). Therefore, it is likely that both oxidative and cell envelope stress can regulate YfiN activity; our biofilm and motility results support the previously reported effects of oxidative stress (66). Moreover, YfiN has been reported to localize to the mid-cell and arrest division in *Salmonella enterica* and *Escherichia coli* and regulate cell division in response to reductive stress (67). Interestingly, increased mid-cell localization was not observed in *P. aeruginosa* (67), indicating that this protein contributes to different functions in different organisms.

The YfiBNR system shows conservation among other Gram-negative microbes beyond *E. coli*, including those organisms known to make biofilms. *Serratia* and *Yersinia*, also in the family Yersiniaceae, utilize the homologous system HmsCDE (38). Like YfiBNR, the HmsCDE system is implicated in biofilm growth and response to redox stress. Deletion of the *yfiN* homolog, *hmsD*, in *Y. pestis* leads to decreases for *in vitro* and *in vivo* biofilm formation (68). Null mutants of *hmsD* exhibit decreased biofilm biomass in reducing environments (69); these previous findings and our work here are consistent with the importance of YfiN in biofilm maintenance, particularly in response to redox conditions. Furthermore, in liquid culture, HmsD and HmsC (YfiR homolog) expression is more than doubled during the late stationary phase, and compared to the control, the Δ*hmsD* mutant exhibits a severe growth defect when grown in media containing H_2_O_2_. We did not observe increased sensitivity of the Δ*yfiN* mutant to H_2_O_2_ in planktonic conditions, again emphasizing the differential functions of this system in different organisms.

## MATERIALS AND METHODS

### Strains and media

UCBPP-PA14 (PA14) was used in all the experiments. *P. aeruginosa* was routinely streaked onto lysogeny broth (LB) plates containing 1.5% agar prior to overnight culturing in LB liquid cultures at 37°C. As needed, LB was supplemented with 10mg/ml gentamicin (Gm) and 50mg/ml kanamycin (Kan). Biofilm medium used in static 96-well crystal violet assays was composed of M63 medium supplemented with 1 mM MgSO4 and 0.4% (wt/vol) L-arginine monochloride. For medium replacement assays, plates were grown as indicated, spent medium discarded and biofilms washed twice prior to the addition of fresh medium. To limit the formation bubbles caused by the addition of H_2_O_2_ during microfluidic-based biofilm assays, we used buffered KA biofilm medium, a modification K10T medium that contains 50mM Tris-HCl (pH 7.4), 0.61mM MgSO4, and 0.4% arginine (70).

### Static biofilm assays and quantification

Overnight cultures were inoculated into 96-well U-bottom polystyrene plates (Costar) containing M63-based biofilm medium and grown for the specified time at 37°C in a hydrated container prior to washing, staining with crystal violet (CV) and solubilization of the CV stain with 30% glacial acetic acid (71). The presence of the biofilm was quantified by measuring the extent of biofilm-associated CV measured in a spectrophotometer to determine the optical density at 500 nm (OD550).

For biofilms imaged microscopically in static assays, overnight cultures were prepared as described above and inoculated into 8-well glass-bottom dishes containing M63 biofilm medium and grown at 37°C for the desired length of time prior to imaging with a Nikon Eclipse Ti inverted microscope where a minimum of 6 fields of view were captured. To assess the viability of biofilms grown in glass bottom dishes, growth medium was discarded, biofilms washed twice with fresh medium, and the biofilms immersed in fresh medium containing propidium iodide 10 min prior to imaging. To assess the concentration of c-di-GMP we utilized P_c*drA*_-*gfp* fusion expressed from a multicopy pMQ72 plasmid (72) which was maintained in overnight cultures supplemented with Gm prior to inoculation into antibiotic-free M63 biofilm medium.

### Microfluidics

Biofilms were visualized under flow in microfluidics chambers kindly provided by the Nadell laboratory at Dartmouth College. Chambers used polydimethylsiloxane (PDMS) bonded to a cover glass (1.5 36 mm 60 mm; Thermo Fisher, Waltham MA) through soft lithography techniques (73, 74). Overnight bacterial cultures were centrifuged, resuspended in biofilm medium, adjusted to an OD600 of 1, pipetted into microfluidics chambers, and allowed to attach for 1 h. Tubing (catalog no. 30 Cole Palmer PTFE) to transport influent and effluent medium was attached first to BD 5-ml syringes containing biofilm medium, then to the microfluidics chambers, and then to syringe pumps (Pico Plus Elite; Harvard Apparatus) operating at a flow rate of 0.75µl/min.

### Image acquisition and data analysis

All microscope images were acquired using Nikon Elements AR driving a Nikon Eclipse Ti inverted microscope and a Hamamatsu ORCA-Flash 4.0 camera. Samples were imaged with either a Plan Apochromat 100 DM Oil or Plan Fluor 40 DIC M N2 objective. Fast scan mode and 2×2 binning were used for imaging. All images were collected in a temperature controlled environmental chamber set to 37°C. Images were processed with background subtraction and signal strength quantified by measuring the mean signal intensity/pixel through the Integrated Density (IntDen) function. To quantify single-cell motility, images were inverted, a threshold applied and the TrackMate program (75) used.

### Statistical analysis

Data were analyzed using GraphPad Prism 8. Unless otherwise noted, data are representative of the results from at least three independent experiments. A Student T-test with a Dunnett’s or Sidak’s multiple comparisons post-test, as indicated, were used to compare results and to assess significance.

### Potential homolog identification and phylogenetic tree construction

The nucleotide sequence of YfiBNR in PAO1 (NC_002516.2:1212717-1215074) was used as a query in a Nucleotide BLAST search (40). Bacterial genomes within the RefSeq Representative genomes database were searched with discontiguous megablast. The expectant threshold was set to 0.0001; all other default algorithm parameters were used. Sequences with >75% query cover were aligned using MAFFT (41). FastTree 2.1.11 was used to construct a phylogenetic tree based on the aligned sequences, using the GTR+CAT model and Gamma20-based likelihoods (42). 1000 re-samples were used to calculate bootstrap support for the tree, and a nonhomologous human protein was used as an outgroup to root the tree. Interactive Tree of Life version 4 was used to visualize the phylogenetic tree.

## Acknowledgements

We thank Carey Nadell for providing microfluidic devices, Zdenek Svindrych for providing imaging guidance, Olga Zhaxybayeva for providing bioinformatic advice, and Thomas Wood for generously and speedily sending strains. We also thank Gillian Kenyon for transformation selected strains with fluorescent marker plasmids. This work was supported by the NIH (R37 AI83256 to GAO) and the James O. Freedman Presidential Scholar Award to SAK.

